# Phylogenomic inference suggests differential deep time phylogenetic signals from nuclear and organellar genomes in gymnosperms

**DOI:** 10.1101/2025.03.16.643507

**Authors:** Yu En Lin, Chung-Shien Wu, Yu-Wei Wu, Shu-Miaw Chaw

## Abstract

The living gymnosperms include about 1,100 species in five major groups: cycads, ginkgo, gnetophytes, Pinaceae (conifers I), and cuppressophytes (conifers II). Molecular phylogenetic studies have yet to reach a unanimously agreed relationship among them. Moreover, cytonuclear phylogenetic incongruence has been repeatedly observed in gymnosperms. We collated a comprehensive data set from available genomes and added our own high-quality assembly of a species from Podocarpaceae (the 2nd largest conifer family) to increase sampling width. We used these data to infer reconciled nuclear species phylogenies using two separate methods to ensure robustness of our conclusions. We also reconstructed organelle phylogenomic trees from 41 mitochondrial and 82 plastid genes. Our nuclear phylogeny consistently recovers the Ginkgo-cycads clade as the first lineage split from other gymnosperm clades and the Pinaceae as sister to gnetophytes (the Gne-pines hypothesis). In contrast, the mitochondrial tree places cycads as the earliest lineage in gymnosperms and gnetophytes as sister to cupressophytes (the Gne-cup hypothesis) while the plastomic tree supports the Ginkgo-cycads clade and Gnetophytes as the sister to Cupressophytes. We also examined the effect of mitochondrial RNA editing sites on the gymnosperm phylogeny by manipulating the nucleotide and amino acid sequences at these sites. Only complete removal of editing sites has an effect on phylogenetic inference, leading to a closer congruence between mitogenomic and nuclear phylogenies. This suggests that RNA editing sites carry a phylogenetic signal with distinct evolutionary traits.

## 1. Introduction

Gymnosperms are one of the two clades of extant seed plants that originated in the Middle-Devonian period about 385 million years ago [1, 2]. In contrast to angiosperms that comprise some 350,000 species, gymnosperms are much less diverse and include only 1,174 species [2]. Nonetheless, gymnosperms dominate huge swaths of the earth’s landmass, especially the conifers (Pinaceae and Cupressaceae) in the Northern Hemisphere and most Araucariaceae and Podocarpaceae genera in the Southern Hemisphere [3]. Consequently, gymnosperms are of great economic and ecological importance in several countries. For example, many Cupressaceae (cypress family), including false cypress (*Chamaecyparis*), China fir (*Cunninghamia*), bald cypress (*Taxodium*), and arborvitae (*Thuja*) are valuable as both timber sources and ornamentals. The yew family (Taxaceae), including about 30 species in six genera, produces taxane compounds useful in anticancer therapies [4].

Advances in sequencing, alignment, and clustering methods starting in the 1990s prompted a reassessment of the classical taxonomic relationships among gymnosperms using molecular data. These analyses categorized gymnosperms into five widely accepted groups: cycads (Cycadales), ginkgo (Ginkgoales), gnetophytes (Gnetales), pine family (Pinales or conifers I), and cuppressophytes (conifers II) [5, 6]. The phylogenetic relationships among these groups are a subject of extensive debate [7], comparable to the controversies surrounding primitive (or early divergent) dicots, monocots, and eudicots. The gymnosperm group monophyly and the placements of ginkgo and gnetophytes were once highly contentious. The initial molecular phylogenetics studies proposed five alternative hypotheses: (1) The extant gymnosperms comprise a monophyletic group; (2) Cycads and ginkgo are sister groups (i.e., the Ginkgo-cycads hypothesis); (3) Gnetophytes are sister to all conifers (gnetifer hypothesis); (4) Gnetophytes are sister to Pinaceae (the Gne-pines hypothesis).; and (5) Gnetophytes are sister to Cupressophytes (the Gne-cup hypothesis) (Figure 1) [5, 6, 8].

**Figure 1.**
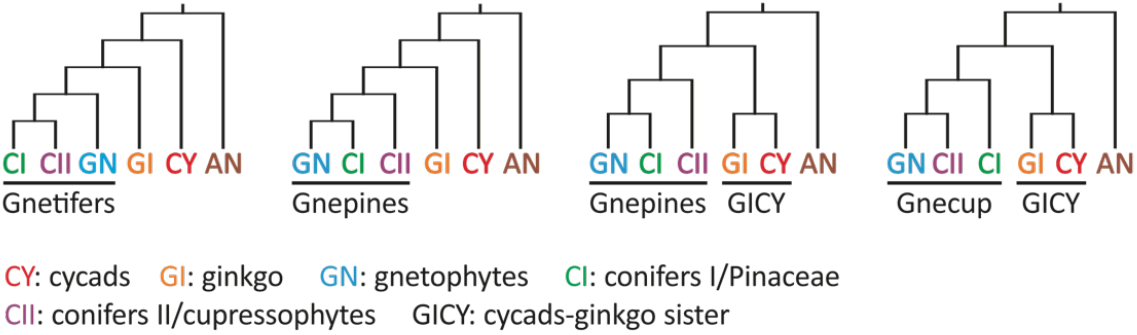
Combinations of the four major hypotheses concerning the phylogenetic relationships among the five extant gymnosperm groups. Angiosperms were the designated sister-group of gymnosperms.

Substantial progress in understanding angiosperm evolution is driven by extensive nuclear phylogenomics studies [9, 10]. Although a few recent studies have analyzed the gymnosperm phylogeny using large transcriptomic datasets [11, 12], this line of investigation remains relatively underdeveloped. Nuclear genome sequencing is challenging in gymnosperms due to their large genomes (ca. 5-28 Gb [13]), with exceptionally long genes, introns, and many repeated sequences [14-18]. Fortunately, the advancements in sequencing technology over the past five years has made whole genome assemblies more feasible and cost effective.

Phylogenomic trees based on whole nuclear genomes are largely inferred from single-copy orthologous genes, excluding the numerous genes including multiple paralogs and thus only accounting for a small, and possibly biased, portion of the phylogenetic signal accumulated in genomes during evolution. Moreover, some phylogenomic trees are constructed from transcriptomes [11, 12]. This can lead to several problems: (1) some transcripts are not in full length, causing sampling bias or alignment errors; (2) transcripts at low levels might lead to inaccurate assembly (or come from DNA contamination [19]); and (3) transcripts represent only a partial set of genes specific to cell types, developmental stages, and growing conditions sampled. To address these issues, we gathered 20 well-assembled seed plant nuclear genomes and incorporated two phylogeny inference and incorporated two phylogeny inference strategies, ASTRAL-Pro2 (ASTRAL for Paralogs and Orthologs) and SpeciesRax [20-22], into this study. These recently developed gene tree-based methods can account for gene duplications, losses, transfers, and to some degree incomplete lineage sorting by reconciling multiple-copy gene families when building species trees.

In addition to nuclear genomes, organelle markers/genomes have been frequently used to resolve plant phylogenetic relationships at different taxonomic levels [5, 6, 23]. As the amount of nuclear and organellar data increases, the number of cases of discordance in phylogenies between nuclear and organellar trees, known as cytonuclear incongruence, has likewise grown (see review in Kao et al. 2022) [24]. For instance, plastomes were widely used in green plant phylogenetics for their small size, predominantly uniparental inheritance (mostly paternal in gymnosperms), stable genomic structure, resistance to recombination, and generally low nucleotide substitution rates [25-34]. However, in gymnosperms, plastid-nuclear incongruence has been consistently observed since the earliest study [35]. These discrepancies contributed, among other things, to the continuing controversy over the placement of ginkgo and gnetophytes [6, 36-43]. To date, studies on gymnosperm plastome phylogeny have unanimously agreed on the ginkgo-cycad sisterhood. Nonetheless, the position of gnetophytes is still debated with earlier studies supporting either the Gne-pines or Gne-cup hypothesis (see review by Chaw et al., 2018) [27].

In contrast to the extensive studies on gymnosperm phylogeny using plastomes, mitogenomic phylogeny has received significantly less attention. This can be best attributed to the highly variable size, complex structure, genome rearrangements, and intergenomic interactions in plant mitogenomes [44-50]. These complicated genomic features render the assembly and syntenic analysis of mitogenomes difficult. As a result, most previous studies only extracted and combined the 40/41 conserved genes in plant mitogenomes for phylogenetic inference (e.g. Liu et al. 2014) [51]. However, since plant mitogenomes have a slow substitution rate, conserved gene numbers, and rearrangement in some gene groups, for example, *nad5-nad4-nad2* [52-54], it was suggested that plant mitogenomes can harbor ancient phylogenetic signals in bryophytes and other basal land plants [52]. Therefore, in this study we consider using mitogenomes to help inform gymnosperm phylogeny reconstruction.

Another remarkable feature of gymnosperm mitogenomes is the prevalence of RNA editing sites [55]. RNA editing in plant organelles can correct mutated codons in a transcript to restore conserved amino acids [56-59]. Since mitogenomes evolve slowly, RNA editing sites may contribute to sequence variation, impacting mitophylogeny estimation and potentially causing cytonuclear incongruence. Consequently, many prior studies have tried to evaluate the impact of RNA editing sites on mitogenome phylogenetic inference. Hiesel et al. (1994) favored the use of cDNA rather than DNA to infer phylogeny, as it takes into account the corrected amino acids sequences [60]. Later, Bowe & dePamphilis (1996) found that RNA editing sites did not affect phylogenetic inference and suggested the use of either (but not both) DNA or cDNA for tree building [61]. However, subsequent comparative phylogenetics analyses argued that RNA editing sites carry important phylogenetic signals and advocated for the use of genomic DNA in mitophylogenomics [62-64]. Nevertheless, some studies restore the conserved amino acids or mitigate the effects of RNA editing by changing RNA editing sites from C back to T or even completely removing them [65-72].

In this study, we surveyed reference-level nuclear genomes of 18 gymnosperm species, including 18 published previously (Table 1; as of July 31, 2024) and reporting the first high-quality draft genome from *Naegia nagii* in the podocarp family (Podocarpaceae, details in Supplementary Information). Podocarpaceae, with 19 genera and 187 species mainly found in the Southern Hemisphere, is the largest and most diverse group in the conifers II clade [73]. Incorporation of the first podocarp genome should improve taxonomic sampling of Cupressophytes and fill in the gaps in our understanding of nuclear genomic variation in gymnosperms. We carried out two gene tree-based phylogenetic inference strategies to reconstruct and compare two nuclear phylogenetic trees, and assess the influence of these inference methods on cytonuclear incongruence. Furthermore, we evaluate the impact of RNA editing on mito-nuclear incongruence and consolidate our previous findings regarding the plastid-nuclear incongruent placements of gnetophytes to shed lights on the underlying causes of phylogenetic discordance across the five gymnosperm clades.

**Table 1.** The available and well-annotated gymnosperm nuclear genomes (last access: July, 2024)

| Taxonomic group | Species | BioProject<br>Accession Number | Assembly level | Size<br>(Gb) |
| --- | --- | --- | --- | --- |
| Conifers I | *Larix kaempferi* | PRJNA587041<sup>N</sup> | Scaffold | 10.9 |
|  | *Picea abies* | PRJEB1822<sup>N</sup> | Scaffold | 12 |
|  | *Picea sitchensis* | PRJNA304257<sup>N</sup> | Scaffold | 20.5 |
|  | *Pinus albicaulis* | PRJNA1034085<sup>N</sup> | Scaffold | 27.6 |
|  | *Pinus lambertiana* | PRJNA174450<sup>N</sup> | Scaffold | 27.6 |
|  | *Pinus taeda* | PRJNA174450<sup>N</sup> | Scaffold | 20.5 |
|  | *Pseudotsuga menziesii* | PRJNA174450<sup>N</sup> | Scaffold | 14.7 |
| Conifers II | *Cryptomeria japonica* | PRJDB13806<sup>N</sup> | Chromosome | 9 |
|  | *Metasequoia glyptostroboides* | PRJCA016596<sup>C</sup> | Chromosome | 8.1 |
|  | *Nageia nagi* * | JBITNC00000<sup>N</sup> | Chromosome | 4.3 |
|  | *Sequoia sempervirens* | PRJNA542879<sup>N</sup> | Scaffold | 26.5 |
|  | *Sequoiadendron giganteum* | PRJNA541481<sup>N</sup> | Chromosome | 8.1 |
|  | *Taxus chinensis* | PRJNA730337<sup>N</sup> | Chromosome | 10.2 |
|  | *Torreya grandis* | PRJNA938254<sup>N</sup> | Scaffold | 19.1 |
| Cycads | *Cycas panzhihuaensis* | PRJNA734434<sup>N</sup> | Chromosome | 10.5 |
| Ginkgo | *Ginkgo biloba* | PRJCA001755<sup>C</sup> | Chromosome | 9.8 |
| Gnetophytes | *Gnetum montanum* | PRJNA339497<sup>N</sup> | Scaffold | 2.1 |
|  | *Welwitschia mirabilis* | PRJCA004995<sup>C</sup> | Chromosome | 6.8 |

\*First time sequenced and reported in this project. <sup>N</sup>: NCBI, <sup>C</sup>: CNCB

## 2. Results

We used nuclear, mitochondrial, and plastid genomic data sets to separately infer species trees. To explore the effect of mitochondrial RNA editing sites on phylogenetic inference, we also gathered a smaller dataset that includes 13 gymnosperm mitogenomes (ME-datasets) where C-to-U editing sites in protein-coding genes were verified using transcriptomics (Table S4). Two basal angiosperms, *Amborella* and *Nymphaea*, were used as the outgroups when inferring phylogenetic trees from each data set.

### 2.1 Multiple-copy nuclear gene families support the Ginkgo-cycads sister-relationship and the Gne-pines hypothesis

We were unable to use single copy orthologs to reconstruct nuclear phylogenetic trees because their number drops drastically with increasing taxon count (Figure S4). We therefore used 10,567 (out of more than 40,000) multi-copy gene families that are common to the 18 sampled gymnosperm species (Table 1). We used SpeciesRax and ASTRAL-Pro2 methods to reconstruct the species phylogeny, obtaining identical results with both (Figure 2).

**Figure 2.**
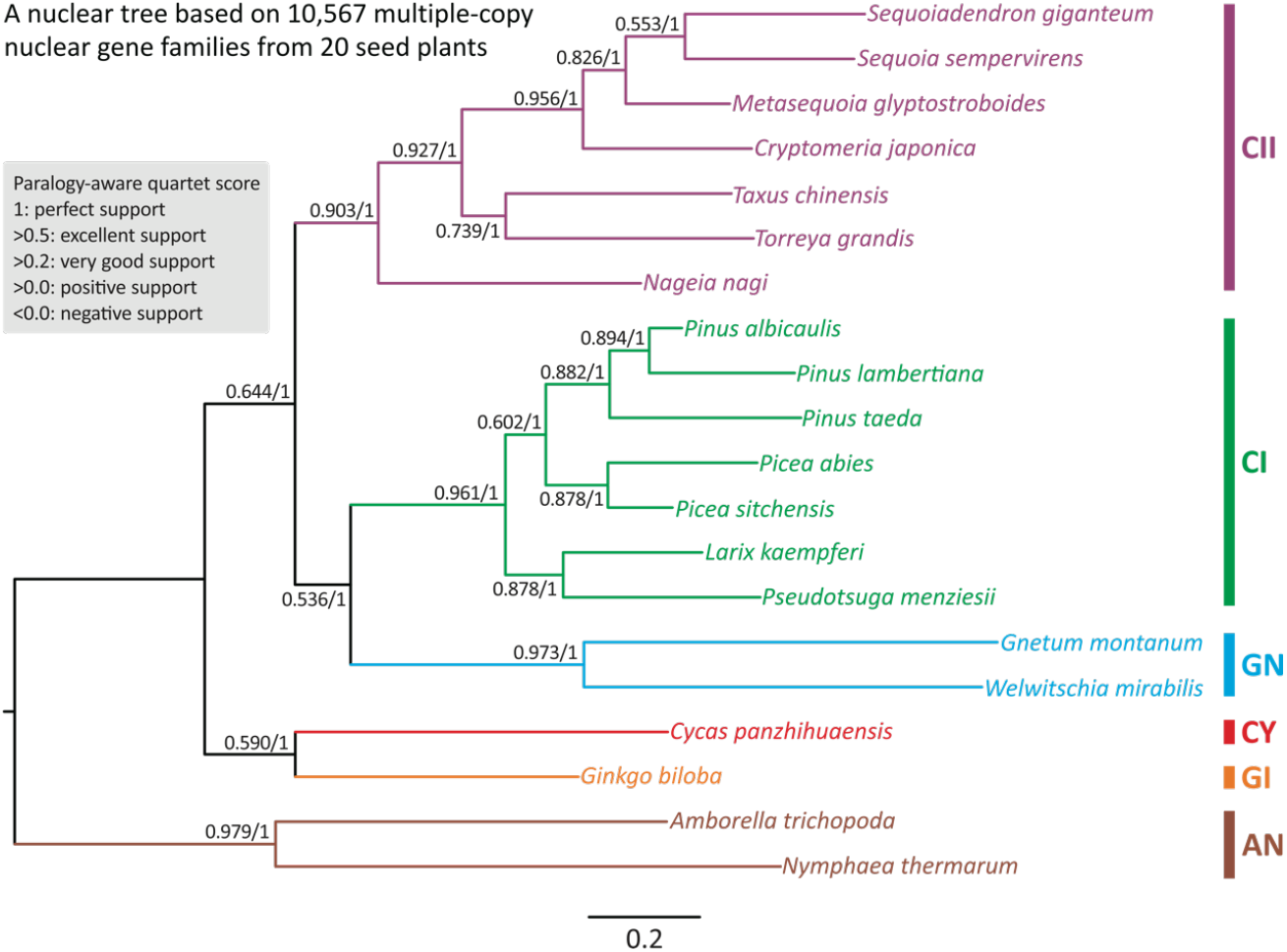
Species trees inferred from 10,567 multiple-copy nuclear gene families across 18 gymnosperms and two angiosperms. The tree framework shown here was constructed using SpeciesRax. Values along branches denote EQPIC scores (before the slashes) and bootstrap support (after the slashes) estimated using SpeciesRax and ASTRAL-Pro2, respectively. AN: angiosperms; GI: ginkgo; CY: cycads; GN: gnetophytes; CI: conifers I; CII: conifers II.

As expected, *Amborella* and *Nymphaea* together form an outgroup clade. At the level of taxonomical orders, our nuclear tree includes two sister clades: the so called “Ginkgo-cycads sister” and “Gne-pines” hypotheses in the gymnosperm phylogeny (Figure 2). SpeciesRax estimates robust extended quadripartition internode certainty (EQP-IC) scores at all nodes, suggesting that the quartets of gene trees also support the two sister clades within the species tree. The strongly supported nodes lead to (1) the Ginkgo-cycads clade (0.59) (2) the three sub-clades: gnetophytes (EQP-IC = 0.973), conifers I (0.961), and conifers II (0.903); and (3) the gnetophytes-conifers I (the Gne-pines subclade) (0.536) clade. Notably, clades (1) and (3) also received 100% bootstrap support in ASTRAL-Pro2 (Figure 2). We thus infer that during gymnosperm evolution (1) the Ginkgo-cycads clade was the first to diverge from the other gymnosperm groups and that (2) Pinaceae (or conifers I clade) is sister to the gnetophytes rather than the conifers II clade. Therefore, the group commonly known as conifers is not monophyletic.

### 2.2 Discordant mitochondrial and plastid phylogenomic trees

The ML tree inferred from both the Mito- and Plastid-datasets recovered cycads, gnetophytes, conifers I, and conifers II each as a monophyletic clade with strong support (all BS > 99%; Figure S5, S6). In addition, the mitochondrial tree topology suggests that (1) cycads diverged first, followed by Ginkgo, and that (2) the “Gne-pines” topology is well resolved (BS = 90%, Figure S5). In stark contrast, our plastid tree resolves the “Gne-cup” topology and the “Ginkgo-cycads” clade with 100% and 98% bootstraps values, respectively (Figure S6). Our results demonstrate that gymnosperm mitochondrial and plastid phylogenies are not only discordant with each other but also differ from the nuclear phylogenomic tree.

### 2.3 Mitochondrial RNA editing sites influence on tree topology

To test if RNA editing sites affect the phylogenetic tree topology, we inferred mitogenomic trees with or without these sites. We needed mitochondrial transcriptome data to find these editing sites. Therefore, the ME-datasets only include 13 gymnosperm species where such data are available. Despite the reduction in taxon count, this set still includes representative species from each of the five extant gymnosperm groups. The RNA editing site numbers vary among genomes, ranging from 99 (*Welwitschia*) to 1,299 (*Keteleeria*) (Figure S7). RNA editing changes some C nucleotides to U in seed plants. It functions as a repair system for maintaining normal mitochondrial protein function, masking some genomic mutations. The sites involved are thus likely under relaxed constraint and might provide misleading phylogenetic signal. Replacing these editing sites with missing data (“N”) or thymine bases (“T”) that reflect their translated sequence would then change the phylogenetic tree topology. However, we do not observe that. Trees inferred from ME-datasets with C-to-N and C-to-T replacements yield identical topologies, both supporting the “Gne-cup” clade, and are the same as the full unaltered data set phylogeny (Figure 3 A-C). However, excluding all RNA editing sites decreases support for the “Gne-pines” clade (BS = 62%) (Figure 3D). Nevertheless, the placement of cycads remains unchanged.

**Figure 3.**
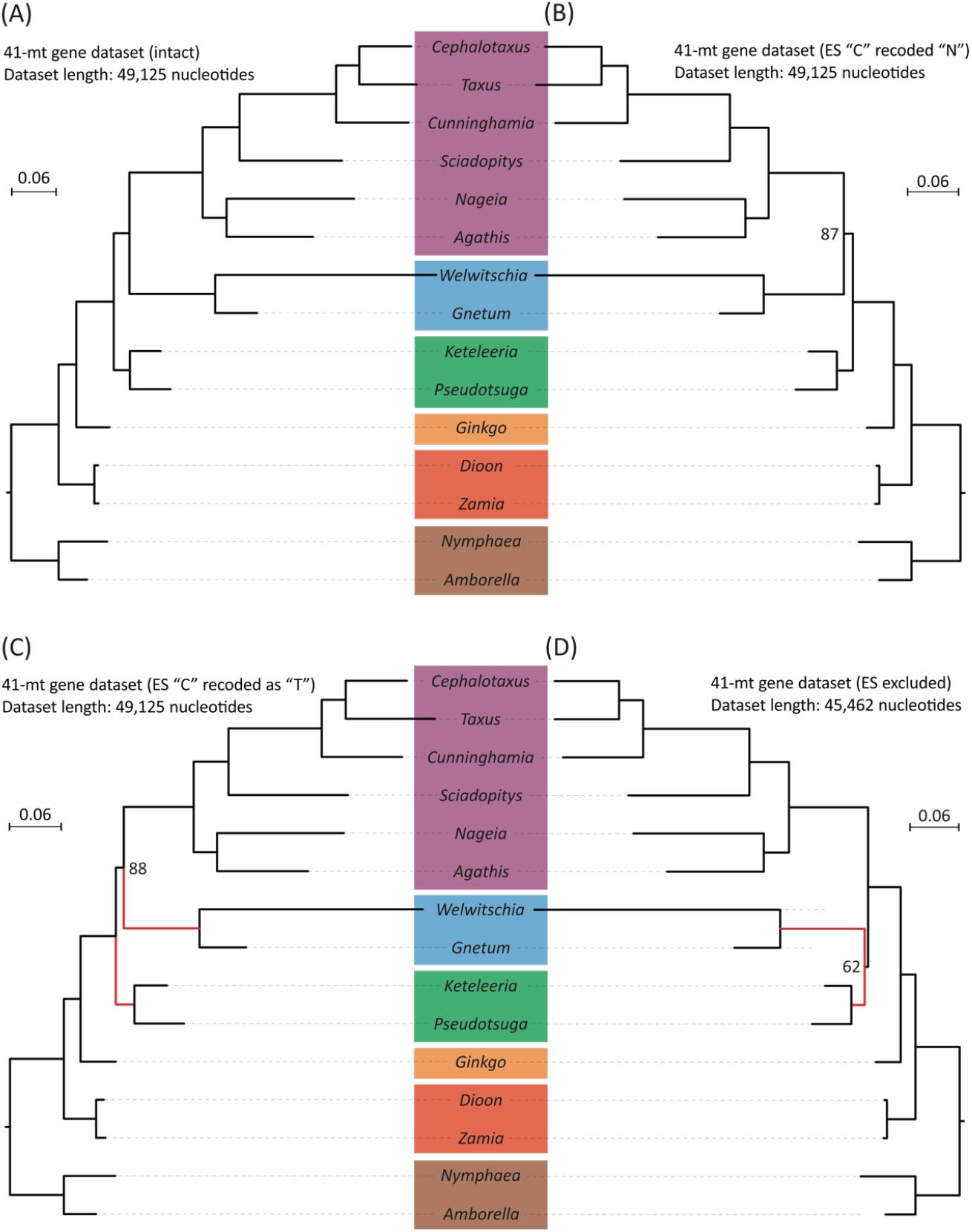
Comparisons of species trees based on four datasets generated from concatenated nucleotide alignments of 41 mt protein-coding genes with and without modifications. (A) A maximum likelihood (ML) tree inferred from the dataset without modification. (B) A ML tree inferred from the dataset where editing sites were replaced with “N” (= missing data). (C) A ML tree inferred from the dataset where edited C nucleotides were recoded as “T” (as in the RNA transcripts). (D) A ML tree inferred from the dataset where alignment positions were removed if they contained and edited nucleotide in any taxon. Bootstrap values are indicated when they are smaller than 100%. The branch length scale bar represents 0.06 substitutions per site. Red lines highlight the placement changes of gnetophytes (here represented by *Welwitschia* and *Gnetum*) between C and D trees.

Because most RNA editing leads to changes in protein sequences, we re-inferred phylogenetic trees using amino acid sequences. We modified the original dataset, in line with the manipulations of the DNA sequences, as follows: (1) amino acids were replaced with “?” if their codons harbored editing sites; (2) all editing sites were recoded as “T” before translation, generating amino acids sequences that are produced *in vivo*; and (3) positions in the alignments were completely excluded if they contained amino acids affected by either synonymous or nonsynonymous editing in any sampled taxon. Phylogenies estimated from the original and the first two modified data sets are the same (Figure 4 A-C). The only difference is that the inference based on the data set without amino acids affected by RNA editing isolates “Gne-pines” into a monophyletic clade with strong support (BS = 94%, Figure 4D). In contrast, the placement of Ginkgo is insensitive to the modifications of sampled taxa numbers and RNA editing sites. All mitochondrial trees, inferred from either nucleotide or amino acids sequences, strongly indicate that Ginkgo diverged after cycads and is sister to the clade comprising gnetophytes, conifers I, and conifers II (Figure S5 and Figures 3, 4; with all BS = 100%).

**Figure 4.**
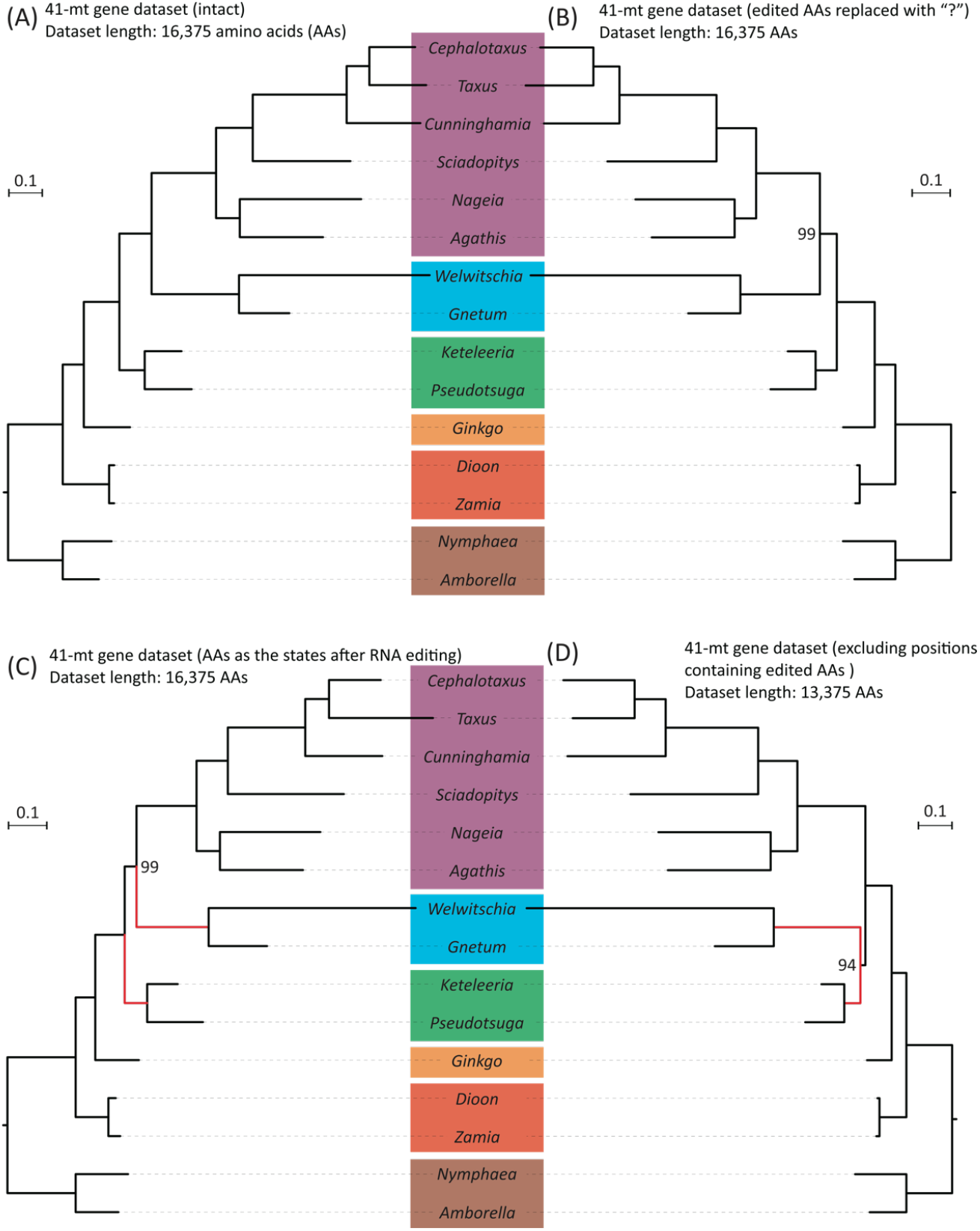
Comparisons of species trees based on four datasets generated from a concatenated amino acid alignment of 41 mt genes with and without modifications. (A) A ML tree inferred from the dataset without modification. (B) A ML tree inferred from the dataset where amino acids affected by RNA editing (including both synonymous and non-synonymous editing) were replaced with “?”. (C) A ML tree inferred from the dataset with amino acids recoded according to the state after RNA editing. (D) A ML tree inferred from the dataset with the aligned positions containing amino acids affected by editing at either synonymous or non-synonymous sites completely removed. Bootstrap values are indicated if they are smaller than 100%. The branch length scale bar represents 0.1 substitutions per site. Red lines highlight the placement changes of gnetophytes (here represented by *Welwitschia* and *Gnetum*) between C and D trees.

In summary, we analyzed three comprehensive datasets, a nuclear and two organelle genomes, of gymnosperms separately to examine the causes of cytonuclear incongruence. Using two separate methods, SpeciesRax and ASTRAL-Pro2, we obtained congruent gymnosperm nuclear phylogenomic trees. ML trees from gymnosperm mitogenomes and plastomes recovered some of the same clades but were not completely in agreement with the nuclear genome-based phylogeny. We also saw that eliminating RNA editing sites from consideration restored “Gen-pines” monophyly. Keeping these sites, regardless of the treatment of their nucleotide state, makes this clade paraphyletic.

## 3. Discussion

Plant nuclear genomes generally mutate at higher rates than the two organelle (or cytoplasmic) genomes [53]. Additionally, while nuclear genomes undergo sexual reproduction and recombination, organelle genomes are generally uniparentally inherited without sexual recombination and thus less genetically variable. The absence of recombination can lead to a build-up of harmful mutations and eventually meltdown of cytoplasmic genomes, a phenomenon known as Muller’s ratchet [74-76], but this process can be slowed down by the low mutation rate in organellar genomes [53].

Interestingly, some plant species (e.g., *Pelargonium, Plantago*, and *Silene*) exhibit extraordinarily accelerated mutation rates in their organellar genomes. In these taxa, biparental inheritance of plastids can occur under mild environmental stress, challenging the long-held belief that organelles are strictly asexual [77]. These discoveries imply that cytonuclear phylogenomic incongruence might be common in plants, as nuclear and organellar genomes follow fundamentally different inheritance patterns. In addition, nuclear genes regulate the function and division of organellar genomes.

Earlier studies attributed the cause of cytonuclear incongruence to a variety of process including long-branch attraction (LBA), incomplete lineage sorting, introgression, gene duplication and loss, distinct organelle inheritance modes, or statistical factors such as sample size [24, 64, 78-83]. In this study, we further scrutinize the causes of cytonuclear incongruence in gymnosperms. To do so, we first constructed a nuclear phylogenomic tree with minimized noise from incomplete lineage sorting, gene duplication and loss, and insufficient sample size. Our objective was to reduce systematic errors in the nuclear phylogenetic inference so that we can justify if the observed incongruence originates from intrinsic evolutionary processes of organellar genomes or from methodological artifacts.

Using the first comprehensive nuclear genome data set to include *Naegia* (Podocarpaceae), we applied two gene tree-based phylogenetic inference methods: ASTRAL-Pro2 and SpeciesRax. Traditional concatenation methods combine per-gene alignments into a single supermatrix to infer a species tree. Since concatenation method only works well with accurate orthology inference [84], it often fails when evolution of genes deviates from that of species due to incomplete lineage sorting or events like duplications, loss, and transfers [85]. On the other hand, gene family tree methods, like ASTRAL-Pro2 and SpeciesRax, preserve the distinct evolutionary history within individual genes family and thus can explicitly account for the underlying processes causing discordance. Specifically, ASTRAL-Pro2 operates under the multispecies coalescent framework, which uses the statistical distribution of gene trees (summarized as quartets) to reduce the influence of incomplete lineage sorting. This method is particularly powerful when multiple-copy nuclear gene families are involved, as it leverages the coalescent process to “average out” the discordance introduced by incomplete lineage sorting. In contrast, SpeciesRax employs a maximum likelihood approach that explicitly models gene duplication, loss, and horizontal transfer events. By incorporating information from both orthologs and paralogs, SpeciesRax is designed to address the complexities of gene family evolution. However, it does not explicitly mitigate the effects of incomplete lineage sorting as the coalescent-based methods do. Together, these two approaches provide a robust alternative to concatenation-based methods by directly modeling the heterogeneous evolutionary processes that shape gene trees [20-22, 86].

Using both methods, we obtained identical topologies of the rooted trees on nuclear data. Our results agree with previous studies that include inferences of rooted as well as unrooted trees [6, 11, 12]. Therefore, nuclear phylogenomics of living gymnosperms have reached a consensus that places Ginkgo and cycads as the earliest-diverging lineage, followed by two sister groups: Gne-pines and cupressophytes (Figure 2). Furthermore, mounting molecular phylogenetic evidence over the past two decades also supports Pinaceae and Gnetophyte monophyly. We thus propose replacing the term “conifers” with Conifers I (Pinaceae) and Conifers II (Cupressophytes) in future research for clarity, since conifers are consistently paraphyletic.

Our analysis of the gymnosperm plastome phylogeny supports the Ginkgo-cycads topology and the Gne-cup topology, in contrast to nuclear-derived Gne-pines topology (Table 2, Figure 2, S6). Numerous factors have been proposed to interpret such incongruence, including incomplete lineage sorting [87], long-branch attraction (LBA) [38, 40, 88, 89], and chloroplast capture [24, 90-94], to name a few. We previously reported significantly accelerated nucleotide substitution rates in gnetophytes, potentially leading to long branch attraction artifacts in phylogenetic reconstruction [38, 40]. Multiple methods that alleviate this artifact restore the Gne-pines topology in plastome trees, suggesting that long branch attraction plays at least a role in mis-inference [38-40, 95]. However, additional analyses would be needed to eliminate or confirm other potential sources of phylogenetic inconsistencies between analyses based on nuclear and organellar data.

**Table 2.**
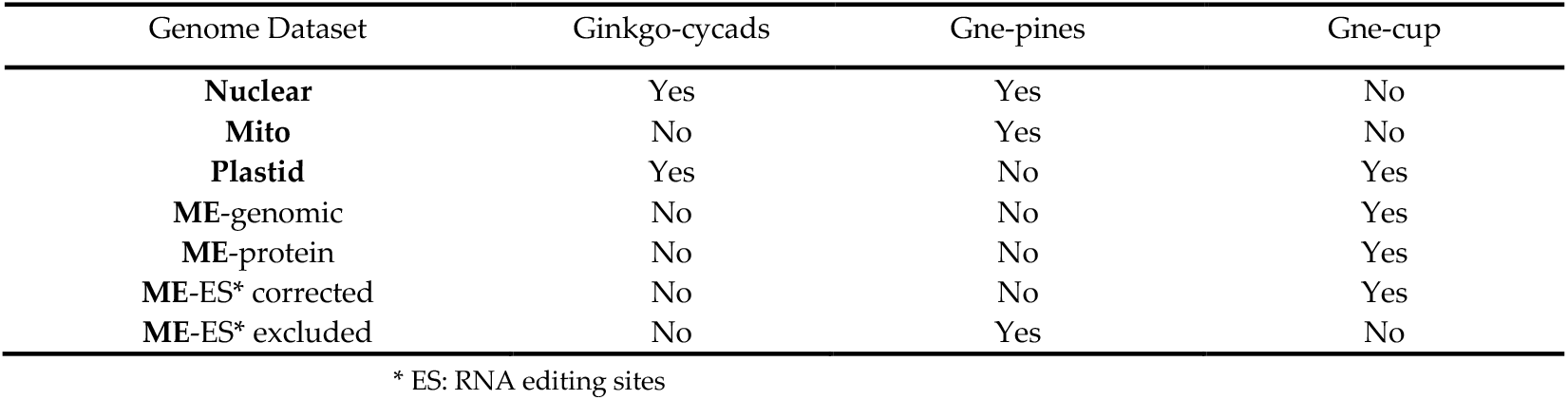
Comparisons of phylogenomic trees inferred from different datasets on controversial clades among the five gymnosperm groups.

| Genome Dataset | Ginkgo-cycads | Gne-pines | Gne-cup |
| --- | --- | --- | --- |
| Nuclear | Yes | Yes | No |
| Mito | No | Yes | No |
| Plastid | Yes | No | Yes |
| ME-genomic | No | No | Yes |
| ME-protein | No | No | Yes |
| ME-ES* corrected | No | No | Yes |
| ME-ES* excluded | No | Yes | No |

\* ES: RNA editing sites

Phylogenies reconstructed from mitochondrial data sets differ from both nuclear- and plastid-based trees. In particular, the mitogenomic phylogeny includes cycads as sister to the remaining gymnosperms (Table 2, Figure 2, S5, S6). Given that RNA editing sites have been proposed as a potential source of phylogenomic incongruence, we further investigated whether modifications of the RNA editing sites at either genomic or protein level could address the observed mito-nuclear discrepancies. Nevertheless, using transcriptomic-verified ME-datasets, our mitogenome species trees show no topological difference before and after the substitution of RNA editing sites (Figure 4A-C), which indicates that the substitution alone is not enough to eliminate the effect of RNA editing, as suggested earlier by other authors [71, 72]. This observation is consistent with the notion that RNA editing sites can influence phylogenetic inference by maintaining accelerated substitution rates and pronounced nucleotide compositional biases, such as an overrepresentation of pyrimidines, even after substitution [55, 59, 64, 71, 72]. In addition, RNA editing frequently produces homoplastic changes, as similar edited states may arise independently in different lineages, thereby introducing convergent noise that confounds the true phylogenetic signal [61]. Moreover, different taxa may exhibit varying efficiencies or patterns of RNA editing. As a result, a uniform substitution does not capture the full extent of the underlying evolutionary variability (Figure S7) [57]. Consequently, the complete removal of RNA editing sites restores the Gne-pines relationship that resembles the nuclear phylogenomic tree topology at both genomic and protein level (Figure 3D, 4D). Furthermore, there is the possibility that the slow substitution rate of mitogenome allow it to carry deep historical signal from ancient events, such as hybridization [67, 68, 82]. This might explain why excluding these sites did not alter the sister relationship of cycads to the remaining gymnosperms (Figure 3D, 4D) [72, 96].

In summary, we suggest that the intrinsic properties of RNA editing sites, like their rapid, biased evolution and convergent behavior, contribute to the observed mito-nuclear incongruence, and that the complete exclusion of them is necessary to mitigate misleading effects. Nonetheless, RNA editing sites in gymnosperms are too few to bias the inference from unedited sites that preserve mitogenomes’ deep and strong phylogenetic signals from ancient hybridization events. We thus see a general preservation of tree topology even after eliminating editing sites, other than the restoration in the placement of gnetophytes [66]. We also note that the “Gne-pines” relationship recovered from the full mitochondrial dataset (Figure S5) was not resolved in the ME-datasets that contains only 13 gymnosperm species (Figure 4A). This suggests that a reduction in taxon sampling can also affect mitochondrial tree topology.

## 4. Materials and Methods

### 4.1 Data access

Nuclear genomes and their annotations were downloaded from the National Center for Biotechnology Information (NCBI, https://www.ncbi.nlm.nih.gov/), the China National Center for Bioinformation (CNCB, https://ngdc.cncb.ac.cn), and the TreeGenes (https://treegenesdb.org/) databases. Incorporating our newly sequenced *Nageia* draft genome (see Supplementary Information), we gathered 18 gymnosperm and two angiosperm genomes that were well-annotated and assembled to scaffold/chromosomal levels (Nu-dataset; Table 1). We used AGAT 1.4.0 (https://github.com/NBISweden/AGAT) to retrieve amino acid sequences of annotated genes from these 20 nuclear genomes. We also gathered amino acid sequences from publicly available mitochondrial and plastid genomes/scaffolds, comprising the Mito- and Plastid-datasets with the former representing 38 (Table S3), and the latter 289 (Table S4) gymnosperm species. Both datasets also include two angiosperms as the outgroups.

### 4.2 Classification of nuclear multiple-copy gene families and construction of gene and species trees

OrthoFinder v3.0.1b1 was used to identify orthologous genes in the 20 sampled nuclear genomes [97]. Only one single-copy and 43,333 multiple-copy nuclear gene families were obtained. To save on computing time, we discarded gene families that are shared by fewer than 10 taxa (i.e. fewer than half of the 20 sampled taxa). This resulted in 10,567 gene families retained for downstream analyses. After using MUSCLE 3.8.1551 to align the sequences [98], the resulting alignments were used to infer gene trees with ParGenes v1.1.2 and its “-m” option for automatically determining the best-fit model [99]. The generated gene trees were pooled before reconciling into species trees using Astral-Pro 2 or SpeciesRax [20-22]

### 4.3 Construction of mitochondrial and plastid trees

The amino acid sequences of mitochondrial and plastid genes were aligned and concatenated into supermetrics using Geneious Prime (www.geneious.com). To better handle sequence heterogeneity in the supermetrics, each gene was designated as a partition to determine the best-fit model. We employed IQtree 2.3.6 for constructing mitochondrial and plastid trees with the “MFP+MERGE” option and 1,000 nonparametric bootstrap replicates [100].

## 5. Conclusions

We present the most comprehensive data set and analysis of gymnosperm cytonuclear incongruence to date. Our results encompass nuclear phylogenomic analyses incorporating, for the first time, a draft Podocarpaceae (from the Southern Hemisphere Conifers II clade) genome. We also construct mitogenome and plastome trees to investigate the factors underlying the observed nuclear-organellar discrepancies. Our extensive analyses suggest that gymnosperm cytonuclear incongruence is likely due to several factors: (1) distinct organelle genome inheritance modes that increase the influence from ancient phylogenetic signals; (2) insufficient taxon sampling; (3) genomic processes, like RNA editing, that can mask the effects of nucleotide substitutions and alleviate functional constraints on protein-coding genes, changing their nucleotide substitution patterns; and (4) Inference bias that stems from incomplete lineage sorting and long branch attraction. Future studies should carefully account for these factors when interpreting discrepancies among inferences based on disparate data sources.

## Supplementary Materials

The following supporting information can be downloaded at: www.mdpi.com/xxx/s1, Figure S1: Flow cytometry of DNA content in *N. nagi*.; Figure S2: Accumulations of scaffolds from the longest to the shortest; Figure S3: Dot-plot analysis of the 13 longest scaffolds.; Figure S4: Variation in the number of identified single-copy gene families with different strategies of taxon sampling.; Figure S5: A maximum likelihood (ML) tree inferred from concatenated amino acid alignments of 42 mitochondrial genes across 41 seed plants.; Figure S6: A maximum likelihood (ML) tree inferred from concatenated amino acid alignments of 82 plastid genes across 288 gymnosperms.; Figure S7: Variation in the total number of RNA editing sites across the 13 sampled gymnosperm mitogenomes.; Table S1: Assembly statistics before and after HiRise scaffolding; Table S2: BUSCO statistics.; Table S3: Gymnosperm plastomes used in this study.; Table S4. Gymnosperm mitogenomes/scaffolds used in this study.

## Author Contributions

S.M.C. conceived and initiated the study. Y.E.L. ad S.M.C. sorted, discussed, wrote, and revised the manuscript. Y.E.L prepared supplementary information and gathered references. C.S.W. carried out phylogenetic tree analyses, made figures, and gave critical comments.

Y.W.W. performed the assembly and annotation of the *Naegia* genome and provided technical support on genomic analyses. S.M.C supervised the experiments and gathered funding.

## Data Availability Statement

All of the raw sequence reads used in this study have been deposited in NCBI under the BioProject accession number JBITNC00000.

## Acknowledgments

We are most grateful for Dr. Bill W. Martin’s encouragement and support for S.M.C. and Y.E.L. to visit his lab for discussing this project, and for Dr. Anthony J. Greenberg’s critical reading and constructive editing of the early version of this manuscript. Special thanks are given to the staff of Dovetail Genomics and Taiwan Genomics for the help and consultation about sequencing the *Naegia* genome and transcriptome.

## Abbreviations

The following abbreviations are used in this manuscript:

NCBI: National Center for Biotechnology Information
CNCB: China National Center for Bioinformation

## References

1. Gerrienne, P., et al., Runcaria, a Middle Devonian seed plant precursor. Science, 2004. 306(5697): p. 856–858.

2. Yang, Y., et al., Recent advances on phylogenomics of gymnosperms and a new classification. Plant Diversity, 2022. 44(4): p. 340–350.

3. Williams, C., Conifer Reproductive Biology. 2009, Springer.

4. Škubník, J., V. Pavlíčková, T. Ruml, and S. Rimpelová, Current perspectives on taxanes: Focus on their bioactivity, delivery and combination therapy. Plants, 2021. 10(3): p. 569.

5. Bowe, L.M., G. Coat, and C.W. DePamphilis, Phylogeny of seed plants based on all three genomic compartments: extant gymnosperms are monophyletic and Gnetales’ closest relatives are conifers. Proceedings of the National Academy of Sciences, 2000. 97(8): p. 4092–4097.

6. Chaw, S.-M., et al., Seed plant phylogeny inferred from all three plant genomes: monophyly of extant gymnosperms and origin of Gnetales from conifers. Proceedings of the National Academy of Sciences, 2000. 97(8): p. 4086–4091.

7. Yang, Y., Z. Yang, and D.K. Ferguson, The Systematics and Evolution of Gymnosperms with an Emphasis on a Few Problematic Taxa. Plants, 2024. 13(16): p. 2196.

8. Nickrent, D.L., C.L. Parkinson, J.D. Palmer, and R.J. Duff, Multigene phylogeny of land plants with special reference to bryophytes and the earliest land plants. Molecular Biology and Evolution, 2000. 17(12): p. 1885–1895.

9. Zuntini, A.R., et al., Phylogenomics and the rise of the angiosperms. Nature, 2024. 629(8013): p. 843–850.

10. Zhang, G. and H. Ma, Nuclear phylogenomics of angiosperms and insights into their relationships and evolution. Journal of Integrative Plant Biology, 2024. 66(3): p. 546–578.

11. Ran, J.-H., T.-T. Shen, M.-M. Wang, and X.-Q. Wang, Phylogenomics resolves the deep phylogeny of seed plants and indicates partial convergent or homoplastic evolution between Gnetales and angiosperms. Proceedings of the Royal Society B, 2018. 285(1881): p. 20181012.

12. Stull, G.W., et al., Gene duplications and phylogenomic conflict underlie major pulses of phenotypic evolution in gymnosperms. Nature Plants, 2021. 7(8): p. 1015–1025.

13. Murray, B.G., Nuclear DNA amounts in gymnosperms. Annals of Botany, 1998. 82(Suppl_1): p. 3–15.

14. Ahuja, M.R. and D.B. Neale, Evolution of genome size in conifers. Silvae Genetica, 2005. 54(1-6): p. 126–137.

15. Morse, A.M., et al., Evolution of genome size and complexity in Pinus. PLoS One, 2009. 4(2): p. e4332.

16. Liu, H., et al., The nearly complete genome of Ginkgo biloba illuminates gymnosperm evolution. Nature Plants, 2021. 7(6): p. 748–756.

17. Wan, T., et al., Evolution of complex genome architecture in gymnosperms. GigaScience, 2022. 11: p. giac078.

18. Zhu, P., T. He, Y. Zheng, and L. Chen, The need for masked genomes in gymnosperms. Frontiers in Plant Science, 2023. 14: p. 1309744.

19. Li, X., P. Zhang, H. Wang, and Y. Yu, Genes expressed at low levels raise false discovery rates in RNA samples contaminated with genomic DNA. BMC Genomics, 2022. 23(1): p. 554.

20. Zhang, C., C. Scornavacca, E.K. Molloy, and S. Mirarab, ASTRAL-Pro: quartet-based species-tree inference despite paralogy. Molecular Biology and Evolution, 2020. 37(11): p. 3292–3307.

21. Zhang, C. and S. Mirarab, ASTRAL-Pro 2: ultrafast species tree reconstruction from multi-copy gene family trees. Bioinformatics, 2022. 38(21): p. 4949–4950.

22. Morel, B., et al., SpeciesRax: a tool for maximum likelihood species tree inference from gene family trees under duplication, transfer, and loss. Molecular Biology and Evolution, 2022. 39(2): p. msab365.

23. Lockwood, J.D., et al., A new phylogeny for the genus Picea from plastid, mitochondrial, and nuclear sequences. Molecular Phylogenetics and Evolution, 2013. 69(3): p. 717–727.

24. Kao, T.T., T.H. Wang, and C. Ku, Rampant nuclear–mitochondrial–plastid phylogenomic discordance in globally distributed calcifying microalgae. New Phytologist, 2022. 235(4): p. 1394–1408.

25. Smith, D.R., Mutation rates in plastid genomes: they are lower than you might think. Genome Biology and Evolution, 2015. 7(5): p. 1227–1234.

26. Xiao-Ming, Z., et al., Inferring the evolutionary mechanism of the chloroplast genome size by comparing whole-chloroplast genome sequences in seed plants. Scientific Reports, 2017. 7(1): p. 1555.

27. Chaw, S.-M., C.-S. Wu, and E. Sudianto, Evolution of gymnosperm plastid genomes, in Advances in Botanical Research. 2018, Elsevier. p. 195–222.

28. Li, H.-T., et al., Plastid phylogenomic insights into relationships of all flowering plant families. BMC Biology, 2021. 19: p. 1–13.

29. Lubna, et al., The dynamic history of gymnosperm plastomes: Insights from structural characterization, comparative analysis, phylogenomics, and time divergence. The Plant Genome, 2021. 14(3): p. e20130.

30. Yang, X., et al., Structural characterization and comparative analysis of the chloroplast genome of Ginkgo biloba and other gymnosperms. Journal of Forestry Research, 2021. 32: p. 765–778.

31. Lian, C., et al., Comparative analysis of chloroplast genomes reveals phylogenetic relationships and intraspecific variation in the medicinal plant Isodon rubescens. PLoS One, 2022. 17(4): p. e0266546.

32. Wang, J., et al., Plant organellar genomes: much done, much more to do. Trends in Plant Science, 2024. 29(7): p. 754–769.

33. Feng, M., et al., The complete plastid genome provides insight into maternal plastid inheritance mode of the living fossil plant Ginkgo biloba. Plant Diversity, 2023. 45(6): p. 752.

34. Shrestha, B., L.E. Gilbert, T.A. Ruhlman, and R.K. Jansen, Clade-specific plastid inheritance patterns including frequent biparental inheritance in passiflora interspecific crosses. International Journal of Molecular Sciences, 2021. 22(5): p. 2278.

35. Palmer, J.D., Comparative organization of chloroplast genomes. 1985.

36. Chaw, S.-M., et al., Molecular phylogeny of extant gymnosperms and seed plant evolution: analysis of nuclear 18S rRNA sequences. Molecular Biology and Evolution, 1997. 14(1): p. 56–68.

37. Palmer, J.D., D.E. Soltis, and M.W. Chase, The plant tree of life: an overview and some points of view. American journal of botany, 2004. 91(10): p. 1437–1445.

38. Wu, C.-S., Y.-N. Wang, S.-M. Liu, and S.-M. Chaw, Chloroplast genome (cpDNA) of Cycas taitungensis and 56 cp protein-coding genes of Gnetum parvifolium: insights into cpDNA evolution and phylogeny of extant seed plants. Molecular Biology and Evolution, 2007. 24(6): p. 1366–1379.

39. Zhong, B., T. Yonezawa, Y. Zhong, and M. Hasegawa, The position of Gnetales among seed plants: overcoming pitfalls of chloroplast phylogenomics. Molecular Biology and Evolution, 2010. 27(12): p. 2855–2863.

40. Wu, C.-S., et al., Comparative chloroplast genomes of Pinaceae: insights into the mechanism of diversified genomic organizations. Genome Biology and Evolution, 2011. 3: p. 309–319.

41. Wu, C.-S., S.-M. Chaw, and Y.-Y. Huang, Chloroplast phylogenomics indicates that Ginkgo biloba is sister to cycads. Genome Biology and Evolution, 2013. 5(1): p. 243–254.

42. Wang, X.-Q. and J.-H. Ran, Evolution and biogeography of gymnosperms. Molecular Phylogenetics and Evolution, 2014. 75: p. 24–40.

43. Ruhfel, B.R., et al., From algae to angiosperms–inferring the phylogeny of green plants (Viridiplantae) from 360 plastid genomes. BMC Evolutionary Biology, 2014. 14: p. 1–27.

44. Palmer, J.D. and L.A. Herbon, Plant mitochondrial DNA evolved rapidly in structure, but slowly in sequence. Journal of Molecular Evolution, 1988. 28: p. 87–97.

45. Chaw, S.-M., et al., The mitochondrial genome of the gymnosperm Cycas taitungensis contains a novel family of short interspersed elements, Bpu sequences, and abundant RNA editing sites. Molecular Biology and Evolution, 2008. 25(3): p. 603–615.

46. Gualberto, J.M., et al., The plant mitochondrial genome: dynamics and maintenance. Biochimie, 2014. 100: p. 107–120.

47. Jackman, S.D., et al., Complete mitochondrial genome of a gymnosperm, Sitka spruce (Picea sitchensis), indicates a complex physical structure. Genome Biology and Evolution, 2020. 12(7): p. 1174–1179.

48. Liu, H., et al., Repetitive elements, sequence turnover and cyto-nuclear gene transfer in Gymnosperm Mitogenomes. Frontiers in Genetics, 2022. 13: p. 867736.

49. Wu, Z.Q., et al., Genomic architectural variation of plant mitochondria—A review of multichromosomal structuring. Journal of Systematics and Evolution, 2022. 60(1): p. 160–168.

50. Xia, C., et al., Complete mitochondrial genome of Thuja sutchuenensis and its implications on evolutionary analysis of complex mitogenome architecture in Cupressaceae. BMC Plant Biology, 2023. 23(1): p. 84.

51. Liu, Y., C.J. Cox, W. Wang, and B. Goffinet, Mitochondrial phylogenomics of early land plants: mitigating the effects of saturation, compositional heterogeneity, and codon-usage bias. Systematic Biology, 2014. 63(6): p. 862–878.

52. Groth-Malonek, M. and V. Knoop, Bryophytes and other basal land plants: the mitochondrial perspective. Taxon, 2005. 54(2): p. 293–297.

53. Drouin, G., H. Daoud, and J. Xia, Relative rates of synonymous substitutions in the mitochondrial, chloroplast and nuclear genomes of seed plants. Molecular Phylogenetics and Evolution, 2008. 49(3): p. 827–831.

54. Mower, J.P., D.B. Sloan, and A.J. Alverson, Plant mitochondrial genome diversity: the genomics revolution. Plant Genome Diversity Volume 1: Plant Genomes, Their Residents, and Their Evolutionary Dynamics, 2012: p. 123–144.

55. Wu, C.S. and S.M. Chaw, Evolution of mitochondrial RNA editing in extant gymnosperms. The Plant Journal, 2022. 111(6): p. 1676–1687.

56. Knoop, V., When you can’t trust the DNA: RNA editing changes transcript sequences. Cellular and Molecular Life Sciences, 2011. 68(4): p. 567–586.

57. Sloan, D.B., Nuclear and mitochondrial RNA editing systems have opposite effects on protein diversity. Biology Letters, 2017. 13(8): p. 20170314.

58. Edera, A.A., C.L. Gandini, and M.V. Sanchez-Puerta, Towards a comprehensive picture of C-to-U RNA editing sites in angiosperm mitochondria. Plant Molecular Biology, 2018. 97: p. 215–231.

59. Dong, S., et al., The amount of RNA editing sites in liverwort organellar genes is correlated with GC content and nuclear PPR protein diversity. Genome Biology and Evolution, 2019. 11(11): p. 3233–3239.

60. Hiesel, R., A. von Haeseler, and A. Brennicke, Plant mitochondrial nucleic acid sequences as a tool for phylogenetic analysis. Proceedings of the National Academy of Sciences, 1994. 91(2): p. 634–638.

61. Bowe, L.M. and C.W. DePamphilis, Effects of RNA editing and gene processing on phylogenetic reconstruction. Molecular Biology and Evolution, 1996. 13(9): p. 1159–1166.

62. Szmidt, A.E., M.-Z. Lu, and X.-R. Wang, Effects of RNA editing on the coxI evolution and phylogeny reconstruction. Euphytica, 2001. 118: p. 9–18.

63. Petersen, G., O. Seberg, J.I. Davis, and D.W. Stevenson, RNA editing and phylogenetic reconstruction in two monocot mitochondrial genes. Taxon, 2006. 55(4): p. 871–886.

64. Picardi, E. and C. Quagliariello, Is plant mitochondrial RNA editing a source of phylogenetic incongruence? An answer from in silico and in vivo data sets. BMC Bioinformatics, 2008. 9: p. 1–11.

65. Bergthorsson, U., K.L. Adams, B. Thomason, and J.D. Palmer, Widespread horizontal transfer of mitochondrial genes in flowering plants. Nature, 2003. 424(6945): p. 197–201.

66. Seberg, O., et al., Phylogeny of the Asparagales based on three plastid and two mitochondrial genes. American Journal of Botany, 2012. 99(5): p. 875–889.

67. Richardson, A.O., et al., The “fossilized” mitochondrial genome of Liriodendron tulipifera: ancestral gene content and order, ancestral editing sites, and extraordinarily low mutation rate. BMC Biology, 2013. 11: p. 1–17.

68. Guo, W., et al., Ginkgo and Welwitschia mitogenomes reveal extreme contrasts in gymnosperm mitochondrial evolution. Molecular Biology and Evolution, 2016. 33(6): p. 1448–1460.

69. Gitzendanner, M.A., et al., Plastid phylogenomic analysis of green plants: a billion years of evolutionary history. American Journal of Botany, 2018. 105(3): p. 291–301.

70. Bell, D., et al., Organellomic data sets confirm a cryptic consensus on (unrooted) land-plant relationships and provide new insights into bryophyte molecular evolution. American Journal of Botany, 2020. 107(1): p. 91–115.

71. Dong, S.S., H.L. Li, B. Goffinet, and Y. Liu, Exploring the impact of RNA editing on mitochondrial phylogenetic analyses in liverworts, an early land plant lineage. Journal of Systematics and Evolution, 2022. 60(1): p. 16–22.

72. Dong, S.-S., X.-P. Zhou, T. Peng, and Y. Liu, Mitochondrial RNA editing sites affect the phylogenetic reconstruction of gymnosperms. Plant Diversity, 2023. 45(4): p. 485.

73. Christenhusz, M.J. and J.W. Byng, The number of known plants species in the world and its annual increase. Phytotaxa, 2016. 261(3): p. 201–217-201–217.

74. Muller, H.J., The relation of recombination to mutational advance. Mutation Research/Fundamental and Molecular Mechanisms of Mutagenesis, 1964. 1(1): p. 2–9.

75. Blanchard, J.L. and M. Lynch, Organellar genes: why do they end up in the nucleus? Trends in Genetics, 2000. 16(7): p. 315–320.

76. Khakhlova, O. and R. Bock, Elimination of deleterious mutations in plastid genomes by gene conversion. The Plant Journal, 2006. 46(1): p. 85–94.

77. Chung, K.P., et al., Control of plastid inheritance by environmental and genetic factors. Nature Plants, 2023. 9(1): p. 68–80.

78. Renoult, J.P., et al., Cyto-nuclear discordance in the phylogeny of Ficus section Galoglychia and host shifts in plant-pollinator associations. BMC Evolutionary Biology, 2009. 9: p. 1–18.

79. Huang, D.I., et al., Whole plastome sequencing reveals deep plastid divergence and cytonuclear discordance between closely related balsam poplars, P opulus balsamifera and P. trichocarpa (S alicaceae). New Phytologist, 2014. 204(3): p. 693–703.

80. Roch, S., M. Nute, and T. Warnow, Long-branch attraction in species tree estimation: inconsistency of partitioned likelihood and topology-based summary methods. Systematic Biology, 2019. 68(2): p. 281–297.

81. Smith, S.A., N. Walker-Hale, J.F. Walker, and J.W. Brown, Phylogenetic conflicts, combinability, and deep phylogenomics in plants. Systematic Biology, 2020. 69(3): p. 579–592.

82. Liu, Y., et al., The Cycas genome and the early evolution of seed plants. Nature Plants, 2022. 8(4): p. 389–401.

83. Duan, L., L. Fu, and H.-F. Chen, Phylogenomic cytonuclear discordance and evolutionary histories of plants and animals. Science China Life Sciences, 2023. 66(12): p. 2946–2948.

84. Altenhoff, A.M., N.M. Glover, and C. Dessimoz, Inferring orthology and paralogy. Evolutionary Genomics: Statistical and Computational Methods, 2019: p. 149–175.

85. Mendes, F.K. and M.W. Hahn, Why concatenation fails near the anomaly zone. Systematic biology, 2018. 67(1): p. 158–169.

86. Yan, Z., et al., Species tree inference methods intended to deal with incomplete lineage sorting are robust to the presence of paralogs. Systematic Biology, 2022. 71(2): p. 367–381.

87. Rose, J.P., et al., Out of sight, out of mind: widespread nuclear and plastid-nuclear discordance in the flowering plant genus Polemonium (Polemoniaceae) suggests widespread historical gene flow despite limited nuclear signal. Systematic Biology, 2021. 70(1): p. 162–180.

88. Bergsten, J., A review of long-branch attraction. Cladistics, 2005. 21(2): p. 163–193.

89. Coiro, M., E.A. Roberts, C.-C. Hofmann, and L.J. Seyfullah, Cutting the long branches: Consilience as a path to unearth the evolutionary history of Gnetales. Frontiers in Ecology and Evolution, 2022. 10: p. 1082639.

90. H Rieseberg, L., J. Whitton, and C. Randal Linder, Molecular marker incongruence in plant hybrid zones and phylogenetic trees. Acta Botanica Neerlandica, 1996. 45(3): p. 243–262.

91. Liston, A., et al., Interspecific phylogenetic analysis enhances intraspecific phylogeographical inference: a case study in Pinus lambertiana. Molecular Ecology, 2007. 16(18): p. 3926–3937.

92. Acosta, M.C. and A.C. Premoli, Evidence of chloroplast capture in south American Nothofagus (subgenus Nothofagus, Nothofagaceae). Molecular Phylogenetics and Evolution, 2010. 54(1): p. 235–242.

93. Nauheimer, L., P.C. Boyce, and S.S. Renner, Giant taro and its relatives: a phylogeny of the large genus Alocasia (Araceae) sheds light on Miocene floristic exchange in the Malesian region. Molecular Phylogenetics and Evolution, 2012. 63(1): p. 43–51.

94. Liu, P.-L., et al., Hedysarum L.(Fabaceae: Hedysareae) is not monophyletic–evidence from phylogenetic analyses based on five nuclear and five plastid sequences. PLoS One, 2017. 12(1): p. e0170596.

95. Zhong, B., et al., Systematic error in seed plant phylogenomics. Genome Biology and Evolution, 2011. 3: p. 1340–1348.

96. Zhong, Z.-R., et al., Maternal inheritance of plastids and mitochondria in Cycas L.(Cycadaceae). Molecular Genetics and Genomics, 2011. 286: p. 411–416.

97. Emms, D.M. and S. Kelly, OrthoFinder: phylogenetic orthology inference for comparative genomics. Genome Biology, 2019. 20: p. 1–14.

98. Edgar, R.C., MUSCLE: multiple sequence alignment with high accuracy and high throughput. Nucleic Acids Research, 2004. 32(5): p. 1792–1797.

99. Morel, B., A.M. Kozlov, and A. Stamatakis, ParGenes: a tool for massively parallel model selection and phylogenetic tree inference on thousands of genes. Bioinformatics, 2019. 35(10): p. 1771–1773.

100. Minh, B.Q., et al., IQ-TREE 2: new models and efficient methods for phylogenetic inference in the genomic era. Molecular Biology and Evolution, 2020. 37(5): p. 1530–1534.

