## Supplementary material for "Phylogenomic inference suggests differential deep time phylogenetic signals from nuclear and organellar genomes in gymnosperms": Supplemantary Information.docx

### Supplementary Information

#### Supplementary Materials and methods

**Flow cytometry assays**

Megagametophytes and young leaves were collected from an about 30 years old *Nageia nagi* tree in Academia Sinica (GPS: 25.0429611,121.6153385). Approximately 50 mg of the plant materials were used in the flow cytometry assays according to Galbraith et al.’s method (1983) [1]. One milliliter of the nuclei suspensions was filtered through a 50-µm nylon filter, followed by staining with 50 µg mL^−1^ propidium iodide. After treatments with 50 µg mL^−1^ RNase, the suspensions were incubated on ice for 10 minutes, and then examined on a Beckman Coulter CytoFLEX flow cytometer.

**DNA and RNA extraction**

*N. nagi*’s megagametophytes were used for DNA preparation using the methods described in Stewart and Via (1993) [2]. DNA libraries were constructed based on the PacBio and Illumina standard protocols and then sequenced on a PacBio Sequel sequencer to generate approximately 38M long reads with the length of 10–25 kb. To yield high-quality RNA, mature seeds were collected from an about 30 years old *Nageia nagi* tree. Seed stratification was conducted in a growth chamber under a 20 °C and dark condition for 20 days. Leaves, stems, and roots were harvested from the 20-day-old seedlings for RNA extraction using the methods described in Kolosova et al. (2004) [3]. After DNase I treatments and polyA tail library construction, the resulted RNA libraries were sequenced on an Illumina NovaSeq 6000 platform to generate 150-bp paired end RNAseq reads.

**Sequencing and genome assembly**

PacBio reads were initially *de-novo* assembled using Canu v2.1.1 with the default parameters [4]. The yielded scaffolds were the templates for Hi-C Scaffolding according to the protocol provided by Dovetail Genomics (<https://dovetailgenomics.com/>). The assembly task was carried out using the HiRise assembler (<https://github.com/DovetailGenomics/HiRise_July2015_GR>). The assembly quality was evaluated by BUSCO systems [5].

**Genome annotation**

Gene prediction was conducted in Braker3 v3.0.3 using both RNA-Seq and protein data as hint inputs. RNA-Seq reads from leaves, stems, and roots were first trimmed using Trimmomatic v0.39 and then mapped to the assembled *Nageia* genome using HISAT2 v2.2.1 [6, 7]. SAMTools v1.9-45 was used to convert SAM files into BAM files, which served as RNA-Seq input for Braker3 [8, 9]. The Orthodb embryophyta v10 FASTA file was downloaded from the OrthoDB database and served as protein input for Braker3 in order to conduct gene prediction [10]. The gene prediction completeness and quality were evaluated using BUSCO v5.7.0 with lineage set to be embryophyta.

#### Supplementary Results

**Flow cytometry of DNA content in *N. nagi***

After several rounds of tests, we found that Tris-MgCl_2_ and CystainTM UV Precise P (Sysmex, Illinois) reagent buffers provided the best condition of nuclei suspensions in cell sorting on flow cytometry assays. Two major picks were observed (Figure S1) in the output. One was detected from megagametophytes, which was about half of the other derived from young leaves. This result agrees with the fact that conifers’ megagametophytes are haploid and their somatic tissues, such as leaves, are diploid in the chromosome numbers. Therefore, conifers’ megagametophytes are here demonstrated to be ideal materials for deciphering their huge genomes. Based on the 1C data, Nageia’s genome size was estimated to be ~5 Gb (Figure S1).

**Chromosomal-level genome assembly**

Table S1 compares the assembled scaffolds before and after Hi-C and HiRise scaffolding. Notably, the N50 value changes from 4,929,268 to 338,021,421 bp, reflecting a great improvement after HiRise scaffolding. This improvement is also noted in Figure S2 where the cumulative length of the scaffolds that reaches the expected genome size is much earlier in HiRise scaffolding than that generated only from PacBio reads. We further assessed the assembly quality with the BUSCO statistics. In Table S2, the statistics of “Complete BUSCOs”, “Complete and single-copy BUSCOs”, “Complete and duplicated BUSCOs”, “Fragmented BUSCOs”, and “Missing BUSCOs” were all improved after HiRise scaffolding. However, these improvements were minor for “Complete BUSCOs” and “Complete and single-copy BUSCOs”. This finding is unprecedented and probably arise from our use of haploid DNA for sequencing and assembly. In contrast, the statistics of “Fragmented BUSCOS” changes from 12 to 9, implying that the HiRise scaffolding is effective in gap closing and contig bridging. The 13 longest scaffolds are amounted to ~4.3 Gb (Figure S3), consistent with both our flow cytometry results and previous chromosomal studies (*N. nagi*: 2n = 26; CCDB, chromosome counts database, accessed date: July, 2022).

We’d report the draft genome sequence of Naegia’s till here. As to the detailed assembly results, predicted genes, and genome syntenic maps with other cuppressophytes will be discussed in a separate report.

#### Supplementary Tables

| Table S1. Assembly statistics before and after HiRise scaffolding | | | | |
| --- | --- | --- | --- | --- |
| Assembly | No. of scaffold | Total length (bp) | N50 | L50 |
| Input assembly | 6,092 | 4,318,943,287 | 4,959,268 | 240 |
| HiRise assembly | 2,283 | 4,319,324,487 | 338,021,421 | 6 |

| Table S2. BUSCO* statistics | | | | |
| --- | --- | --- | --- | --- |
| Assembly | Complete BUSCOs | Complete and single-copy BUSCOs | Complete and duplicated BUSCOs | Fragmented BUSCOs |
| Input assembly | 217 (85.10%) | 205 (80.39%) | 12 | 12 |
| HiRise assembly | 221 (86.67%) | 211 (82.75%) | 10 | 9 |
| *BUSCO version: 4.0.5; Lineage dataset: eukaryota_odb10 | | | | |

#### Supplementary Figures

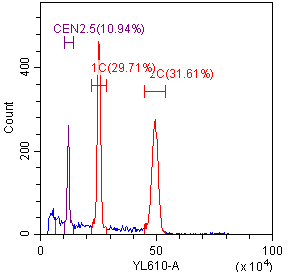

Figure S1. Flow cytometry of DNA content in *N. nagi*. 1C: megagametophyte; 2C: young leaf; CEN: Chicken erythrocyte (control). The estimated 1C-value of *N. nagi* is 5.02–5.12 Gb.

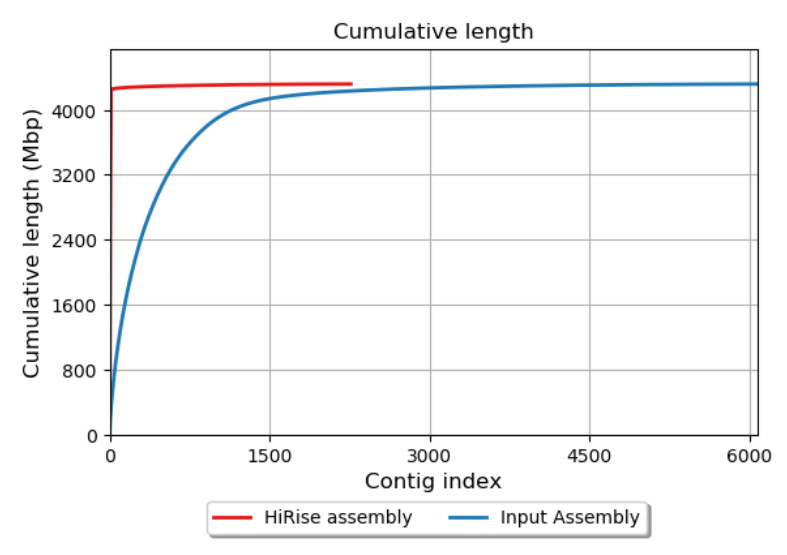

Figure S2. Accumulations of scaffolds from the longest to the shortest

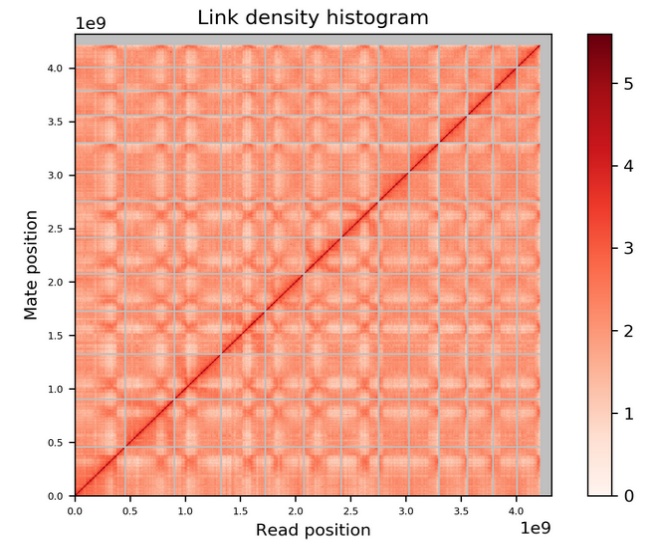

Figure S3. Dot-plot analysis of the 13 longest scaffolds.

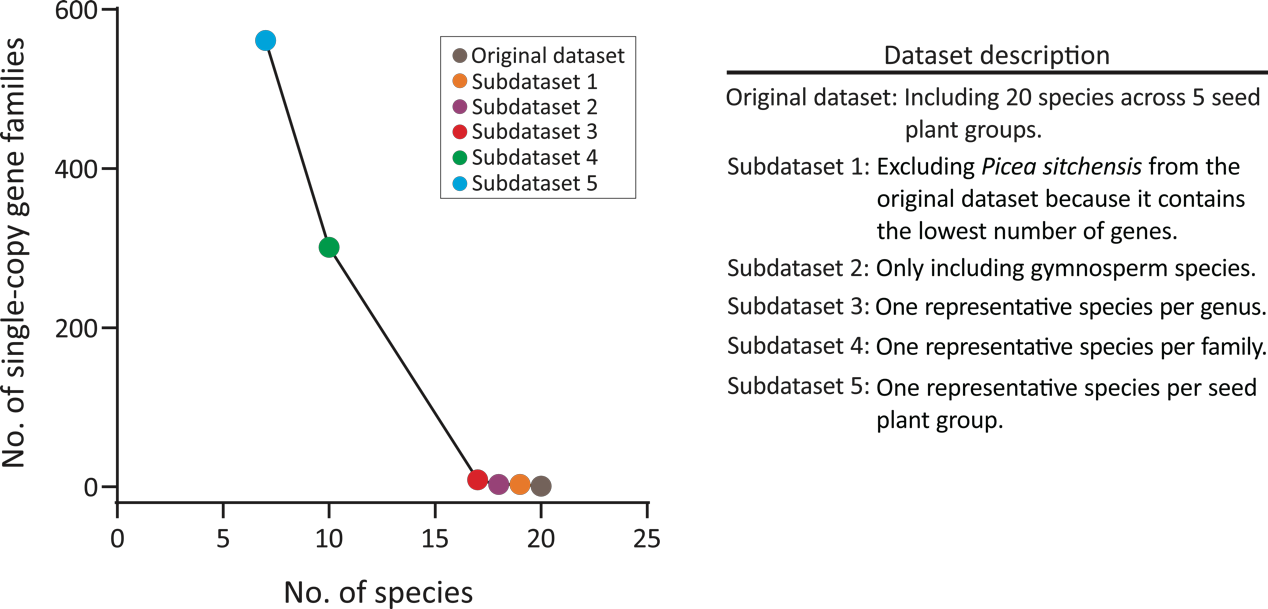

Figure S4. Variation in the number of identified single-copy gene families with different strategies of taxon sampling.

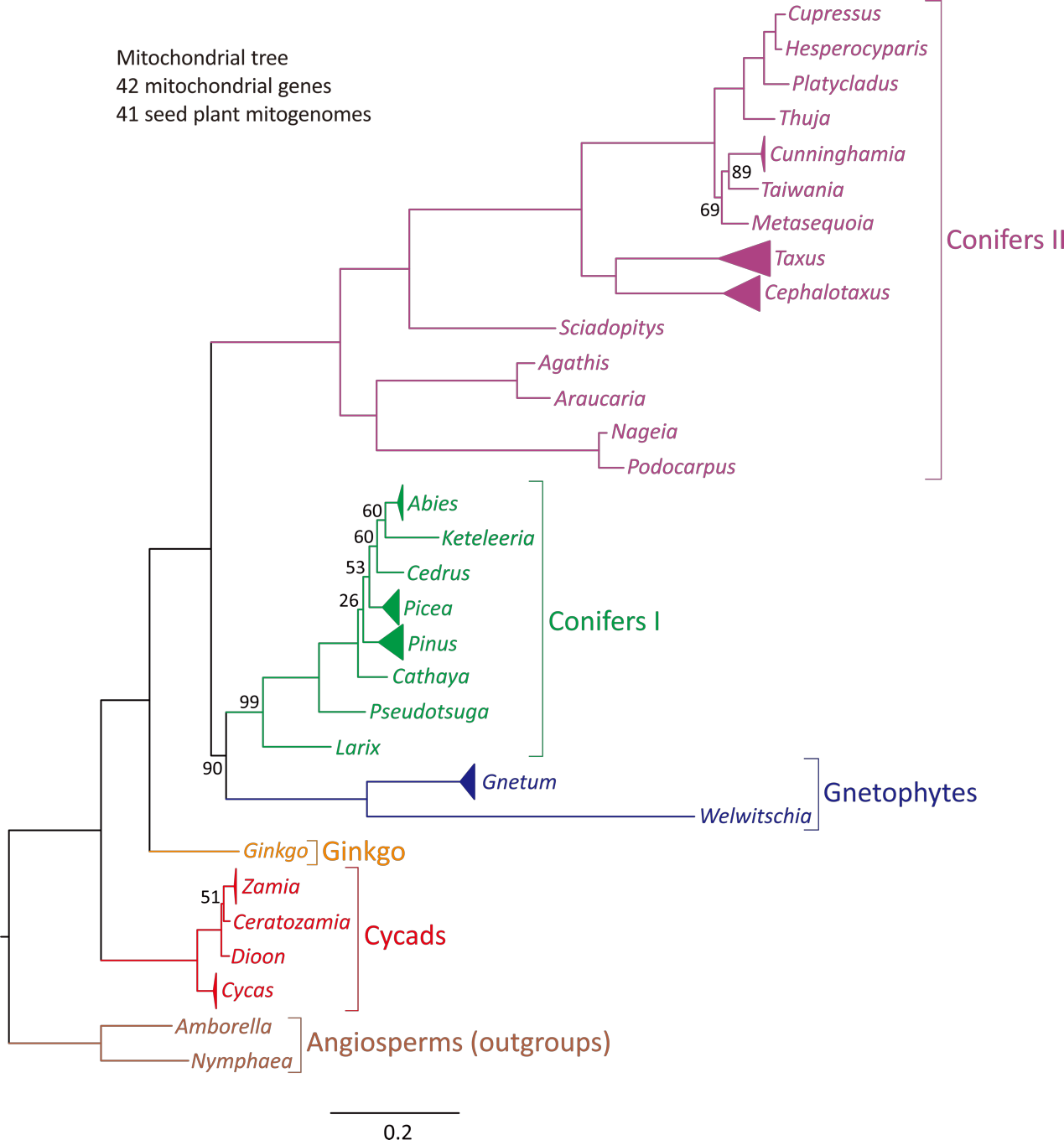

Figure S5. A maximum likelihood (ML) tree inferred from concatenated amino acid alignments of 42 mitochondrial genes across 41 seed plants. Two angiosperms, *Amborella* and *Nymphaea*, were set as the outgroup. Bootstrap values are shown only if they are smaller than 100%

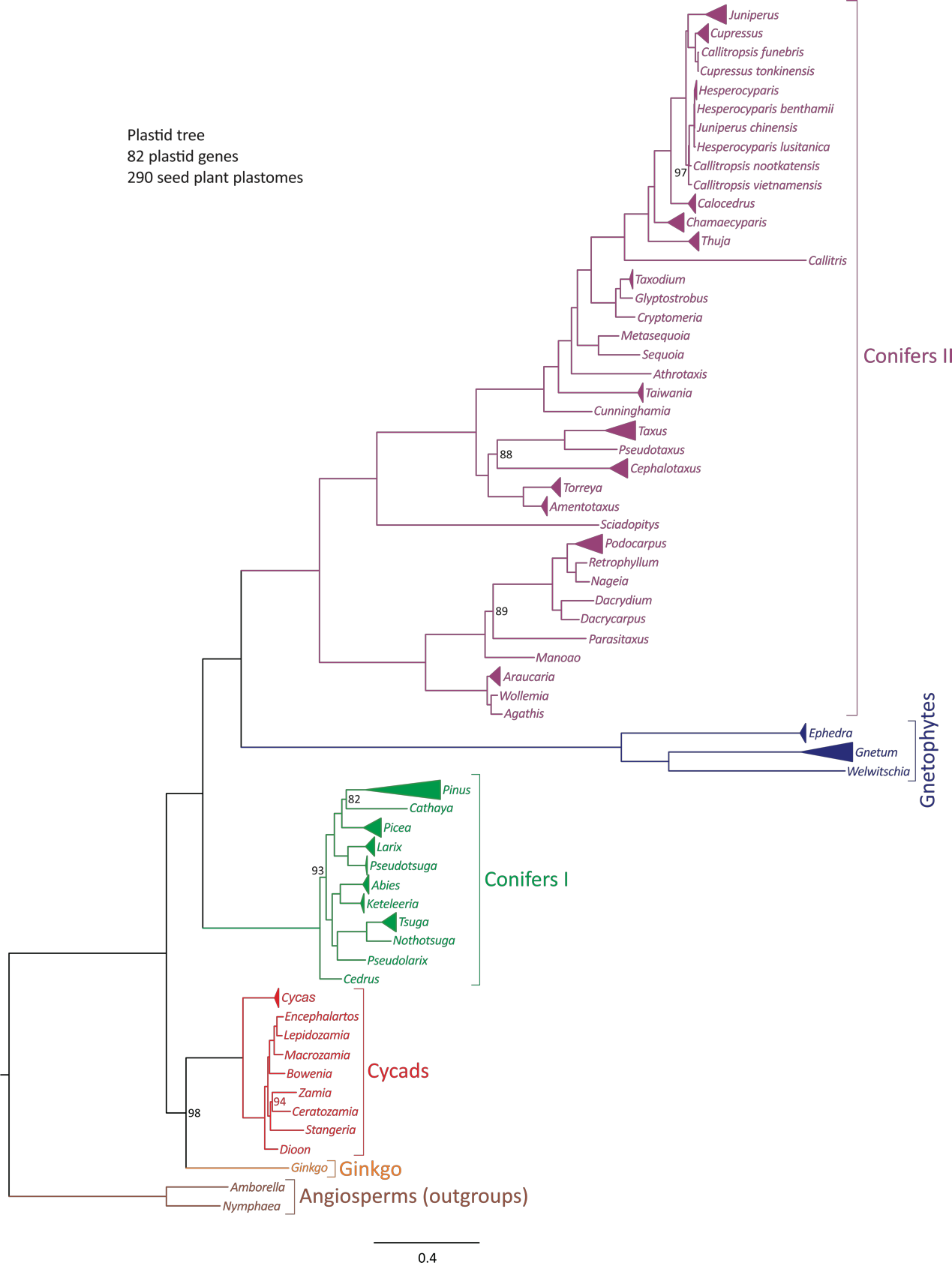

Figure S6. A maximum likelihood (ML) tree inferred from concatenated amino acid alignments of 82 plastid genes across 288 gymnosperms. Two angiosperms, *Amborella* and *Nymphaea*, were set as the outgroup. Bootstrap values are shown only if they are smaller than 100%.

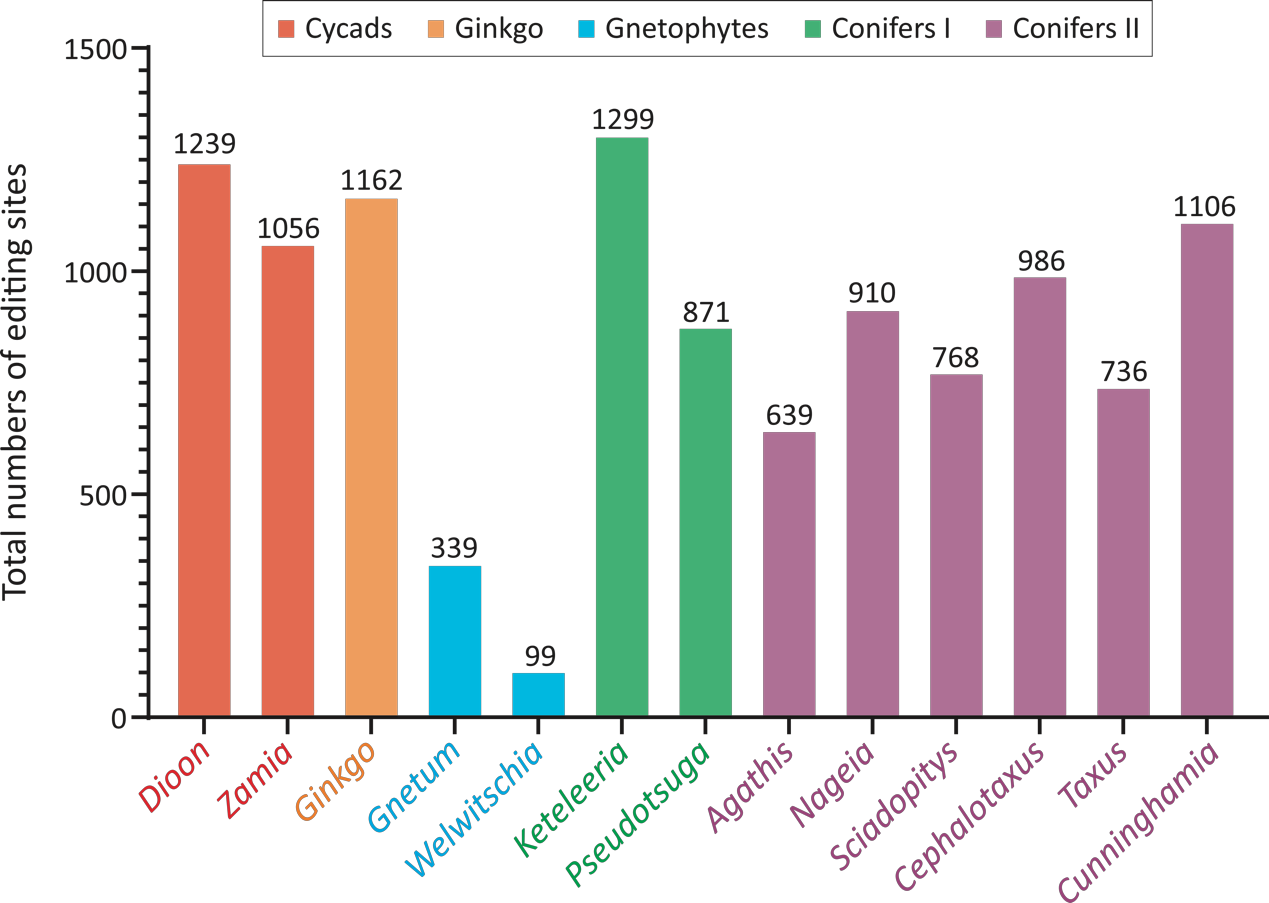

Figure S7. Variation in the total number of RNA editing sites across the 13 sampled gymnosperm mitogenomes.

#### References

1.	Galbraith, D.W., et al., *Rapid flow cytometric analysis of the cell cycle in intact plant tissues.* Science, 1983. **220**(4601): p. 1049-1051.

2.	Stewart, C., Jr and L.E. Via, *A rapid CTAB DNA isolation technique useful for RAPD fingerprinting and other PCR applications.* 1993.

3.	Kolosova, N., et al., *Isolation of high-quality RNA from gymnosperm and angiosperm trees.* Biotechniques, 2004. **36**(5): p. 821-824.

4.	Koren, S., et al., *Canu: scalable and accurate long-read assembly via adaptive k-mer weighting and repeat separation.* Genome Research, 2017. **27**(5): p. 722-736.

5.	Simão, F.A., et al., *BUSCO: assessing genome assembly and annotation completeness with single-copy orthologs.* Bioinformatics, 2015. **31**(19): p. 3210-3212.

6.	Bolger, A.M., M. Lohse, and B. Usadel, *Trimmomatic: a flexible trimmer for Illumina sequence data.* Bioinformatics, 2014. **30**(15): p. 2114-2120.

7.	Zhang, Y., et al., *Rapid and accurate alignment of nucleotide conversion sequencing reads with HISAT-3N.* Genome Research, 2021. **31**(7): p. 1290-1295.

8.	Danecek, P., et al., *Twelve years of SAMtools and BCFtools.* Gigascience, 2021. **10**(2): p. giab008.

9.	Gabriel, L., et al., *BRAKER3: Fully automated genome annotation using RNA-seq and protein evidence with GeneMark-ETP, AUGUSTUS, and TSEBRA.* Genome Research, 2024. **34**(5): p. 769-777.

10.	Kuznetsov, D., et al., *OrthoDB v11: annotation of orthologs in the widest sampling of organismal diversity.* Nucleic acids research, 2023. **51**(D1): p. D445-D451.
